# Spatial transcriptomic landscape of the *Ciona* adult brain: functional zonalisation and cellular composition in a sessile chordate brain and a novel insight into neural gland function

**DOI:** 10.64898/2026.04.13.718160

**Authors:** Xin Zeng, Fuki Gyoja, Ayana Maruo, Nanako Okawa, Ken-ichi Mizutani, Yutaka Suzuki, Kenta Nakai, Takehiro G. Kusakabe

**Affiliations:** Human Genome Center, The Institute of Medical Science, The University of Tokyo, Tokyo, 108-8639, Japan; Department of Computational Biology and Medical Sciences, The University of Tokyo, Kashiwa 277-8563, Japan; Simons Center for Quantitative Biology, Cold Spring Harbor Laboratory, Cold Spring Harbor, 11724, New York, US; Institute for Integrative Neurobiology and Department of Biology, Konan University, Kobe, 658-8501, Japan; Graduate School of Pharmaceutical Sciences, Kobe Gakuin University, Kobe, 650-8586, Japan

## Abstract

The ascidian *Ciona* provides a key model for understanding the evolutionary origin of the vertebrate brain. While the larval nervous system has been extensively characterized, the molecular and cellular organization of the adult neural complex remains poorly defined. Here, we generated spatial transcriptomic maps of the adult *Ciona* neural complex from three individuals, with four serial sections per donor, using the 10x Visium platform. Clustering-based analysis identified five major tissue domains, including the cerebral ganglion, neural gland, ciliated funnel, neural gland duct/dorsal strand, and body wall muscle. To further refine spatial resolution, we computationally reconstructed approximately 980 super-resolution gene expression maps by integrating transcriptomic measurements with histological image features. The super-resolution maps enabled precise delineation of molecular territories within the neural complex. In the cerebral ganglion, high-resolution reconstruction revealed clear molecular zonation, distinguishing the cortex and medulla. Within the cortex, the central region facing the neural gland and anteroposterior distal regions showed distinct molecular properties. In the neural gland, we identified coordinated enrichment of cell-cell interaction- and extracellular matrix–related genes, suggesting specialized structural and physiological properties. We propose that the neural gland play a pivotal role for the cerebral ganglion in maintaining homeostasis, supporting development, and providing a signaling interface, which is reminiscent of a primitive form of the choroid plexus and meninges found in vertebrates. Together, this study provides the first spatially resolved transcriptomic atlas of the adult *Ciona* neural complex and establishes a molecular framework for investigating functional regionalization and brain evolution in chordates.

## INTRODUCTION

The central nervous system (CNS) of the ascidian *Ciona*, one of the closest invertebrate relatives of vertebrates, provides an important model for understanding the evolution of the vertebrate brain ^1^ . For instance, a recent comparative study between *Ciona* larval brain and mouse hypothalamus has provided insights into the evolutionary origin of vertebrate hypothalamic neuroendocrine systems ^2^. Although the larval nervous system of *Ciona* has been extensively characterized at the transcriptomic and cellular levels ^1,3–7^, the adult brain, which is referred to as the neural complex, has received comparatively less attention. One reason for this is technical. While it is relatively easy to analyse gene expression and function at the single-cell level in the brain of ascidian larvae using transient transgenic techniques or whole-mount methods, it is difficult to apply transgenic techniques or perform whole-mount analysis in the brain of adult ascidians, which take several weeks to reach maturity after metamorphosis.

The *Ciona* neural complex consists primarily of the cerebral ganglion and the neural gland and is thought to perform integrative and neuroendocrine-like functions ^8^. Cell lineage tracing studies have revealed that the cerebral ganglion develops during metamorphosis from the neural progenitor cells in the larval brain ^9,10^. However, processes and mechanisms of the neural complex development are still largely unknown. This is primarily due to a lack of information regarding the types of cells that constitute the neural complex and their distribution. Recent anatomical mapping studies have described the peripheral innervation architecture of the adult nervous system ^11^, and neuropeptidomic imaging has revealed regional neurochemical organization within the cerebral ganglion ^12^. However, a spatially resolved transcriptomic map linking cellular identity, molecular programs, and anatomical structure across the adult neural complex is still lacking. Recent advances in spatial transcriptomics enable genome-wide measurement of gene expression while preserving tissue architecture ^13^. In addition, computational approaches that integrate histological image features with transcriptomic measurements can further refine spatial boundaries and reveal substructures beyond the native resolution ^14^. These advances provide an opportunity to systematically characterize the spatial molecular architecture of the neural complex.

Here, we generated spatial transcriptomic data for the adult *Ciona* neural complex and applied computational methods to improve spatial resolution. The resulting high-resolution spatial maps enabled systematic identification of spatially restricted marker genes across the neural complex. We further identified marker genes for the neural gland and three subregions of the cerebral ganglion.

## RESULT

### 1. Spatial transcriptomic landscape of the adult brain of *Ciona*

The adult *Ciona* brain is located in the anterior region of the animal and is mainly composed of two major structures: the cerebral ganglion and the neural gland (**Fig. 1A**). We collected brain sections from three adult *Ciona* individuals using a balanced spatial replicate design (**Fig. 1B**). For each individual, four serial sections from a comparable anatomical region were obtained and distributed across different Visium slides. Histological images acquired during the Visium workflow allowed clear identification of major anatomical features within the brain sections, including the neural gland and cerebral ganglion (**Fig. 1C**). Spatial gene expression profiling of these sections was performed using the 10x Genomics Visium platform. The resulting Visium libraries produced approximately 3.9□×□10^8 to 4.4□×□10^8 reads per sample, with 414–532 spots detected under tissue. Across samples, the median UMI counts per spot ranged from ∼12,000 to ∼20,000, and a median of ∼3,300–4,200 genes were detected per spot (**Supplementary Fig. 1A**). In total, 1,949 spots were retained for downstream analyses.

**Figure 1.**
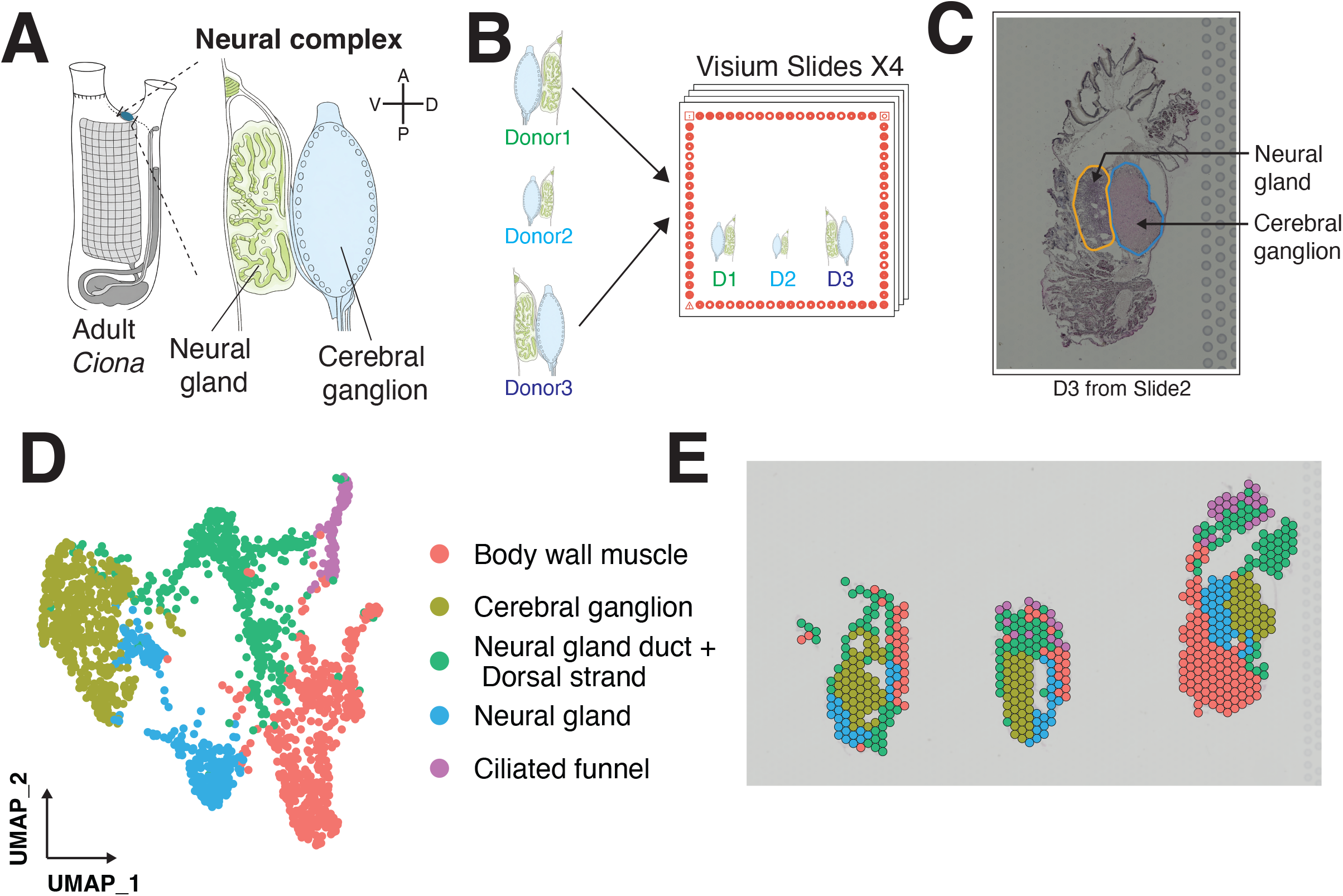
Spatial transcriptomic landscape of the adult Ciona brain. **A** Schematic illustration of the adult *Ciona* neural complex (adapted from ^11^), highlighting the two main brain structures, the cerebral ganglion and the neural gland. Orientation is indicated by anterior–posterior (A–P) and dorsal–ventral (D–V) axes. **B** Experimental design for spatial transcriptomics. Brain tissues from three adult individuals were sectioned serially, with four sections per individual distributed across Visium slides. **C** Representative histological image from the Visium workflow showing anatomical identification of the cerebral ganglion and neural gland. The example shown is donor 3, a section from slide 2. **D** UMAP plots of all transcriptomic spots colored by annotated tissues. **E** Spatial mapping of cell types onto tissue sections.

We performed downstream using Seurat v4 ^15^. To reduce technical variation across samples and individuals, we applied canonical correlation analysis (CCA) to integrate Visium datasets prior to clustering. After batch correction, spots from different samples and individuals were well mixed in the integrated low-dimensional space, indicating effective removal of batch effects (**Supplementary Figs. 1B and 1C**). We then performed graph-based community detection on the integrated data and identified five clusters at the selected resolution (**Fig. 1D**, see Methods). Similar clustering patterns were reproducibly observed across individuals and Visium slides, indicating that the identified clusters were not driven by individual-specific effects (**Fig. 1E and Supplementary Figs. 2A**).

### 2. Tissue annotation for the adult brain of *Ciona*

We assigned each cluster to anatomical structures in the adult *Ciona* brain by combining histological observations and differentially expressed genes. Specifically, we identified marker genes for each spatial cluster based on differential expression analysis (see Methods), which showed clear cluster-specific expression patterns (**Fig. 2A**). Clusters corresponding to the cerebral ganglion and neural gland were identified based on their spatial correspondence to these structures in histological images across sections. For clusters whose anatomical identity could not be unambiguously determined from histology alone, we relied on complementary lines of evidence. We annotated body wall muscle based on strong enrichment of Gene Ontology (GO) terms related to muscle structure and contraction among its marker genes (**Fig. 2B**). In addition, we annotated the ciliated funnel by its anterior-most position and the enrichment of GO terms associated with ciliary organization and motile cilia. Furthermore, we annotated the neural gland duct together with the dorsal strand, based on its reproducible spatial positioning relative to the cerebral ganglion and neural gland across individuals and Visium slides.

**Figure 2.**
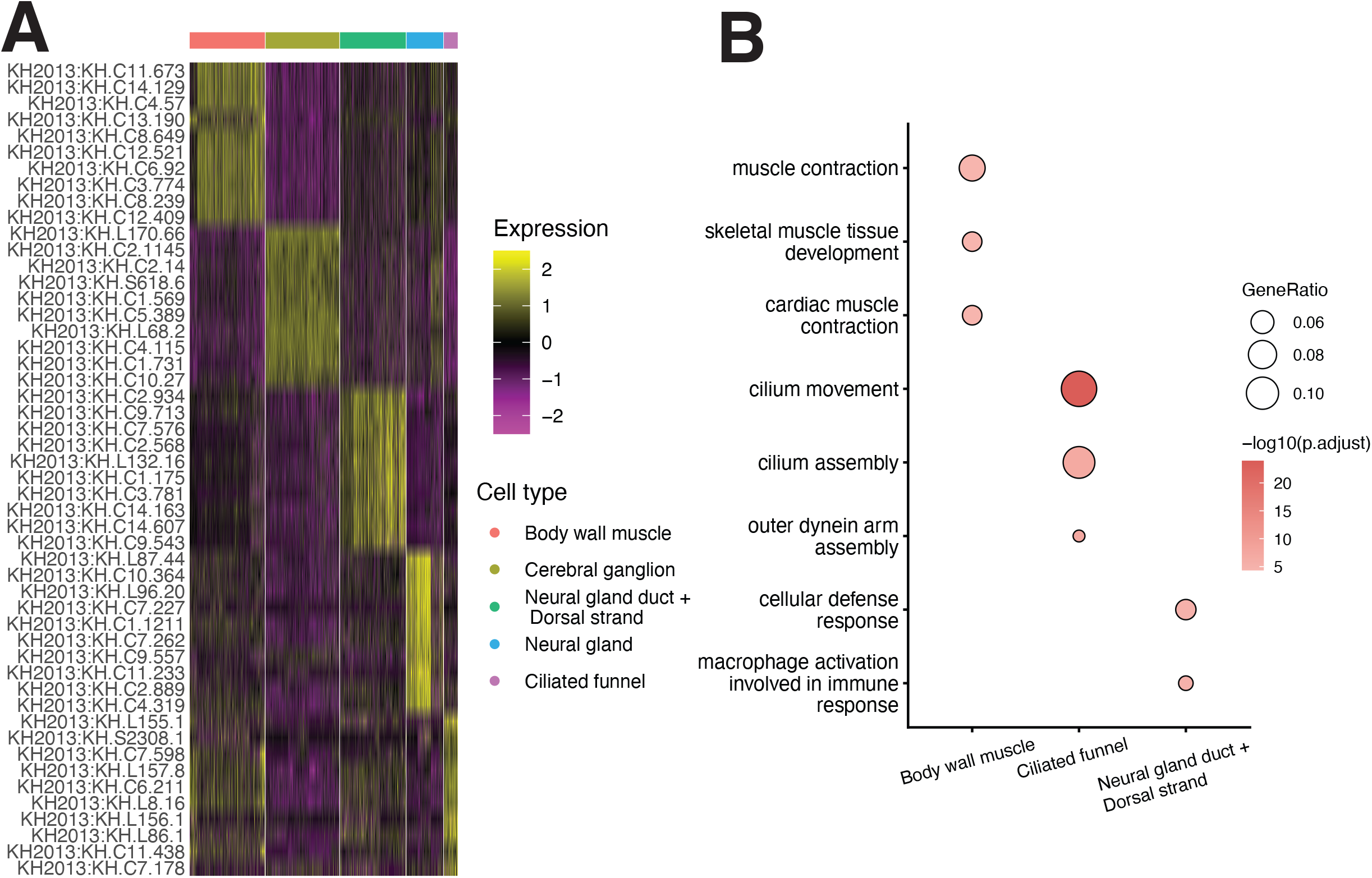
Annotation of clusters in the adult brain of *Ciona*. **A** Heatmap showing scaled expression levels of representative marker genes across all tissue types. **B** Gene Ontology (GO) enrichment analysis of marker genes for body wall muscle, ciliated funnel, and neural gland duct/dorsal strand.

### 3. Construction of super-resolution gene maps of the *Ciona* brain

Although the spatial clusters captured major anatomical structures of the *Ciona* brain, the resolution of Visium transcriptomics data alone limited our ability to resolve finer substructures. To overcome this limitation, we integrated transcriptomic information with spatial image features derived from histological sections using Xfuse ^14^. Compared with raw Visium spot-level expression maps, Xfuse-inferred gene expression maps displayed sharper spatial boundaries and more refined internal organization, enabling clearer delineation of anatomical domains (**Fig. 3A**). In addition, these refined spatial patterns were independently supported by previously published in situ hybridization data ^8,16^, in which the localization of *ci-galp* (KH.S618.6), *ci-ntlp-A* (KH.C2.14), *ci-vfyl/l* (KH.C2.1145), *ci-lf* (KH.C10.27), and *vasopressin receptor* (KH.C9.885) signals consistently showed enrichment in the cortical region of the cerebral ganglion and that of *somatostatin receptor* (KH.C8.337) confined to the neural gland, matching the domains identified by spatial transcriptomics (**Supplementary Figs. 3A-3C**). We then systematically generated super-resolution gene expression maps for all cell-type–specific differentially expressed genes using Xfuse, resulting in approximately 980 high-resolution expression maps in total (**Fig. 3B**).

**Figure 3.**
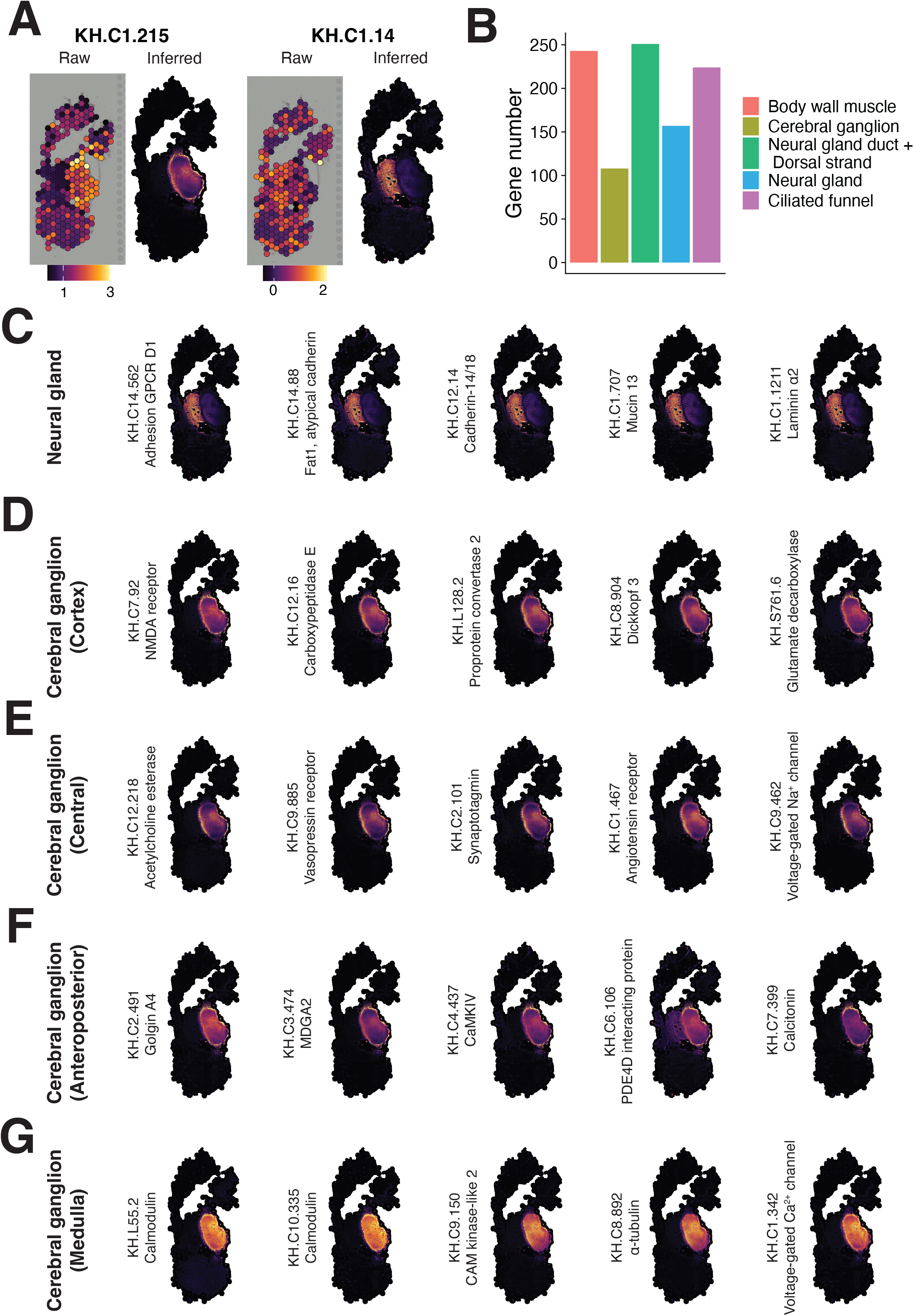
Super-resolution spatial gene maps reveal region-specific marker genes in the adult *Ciona* neural complex. **A** Representative examples of raw Visium spot-level expression maps and corresponding Xfuse-inferred super-resolution maps for selected genes. **B** Number of genes generated by Xfuse for each major anatomical cluster. **C-G** Super-resolution expression maps of representative marker genes enriched in the neural gland (**C**) and different regions of the neural gland (pan-cortex, **D**; central cortex, **E**; anteroposterior cortex, **F**; medulla, **G**).

### 4. Identification of new marker genes for the neural gland

Using super-resolution gene expression maps, we identified a set of new marker genes with spatially restricted expression patterns for the neural complex. In the neural gland, 85 genes showed highly consistent and confined enrichment within this region across individuals (**Fig. 3C; Supplementary Table S1)**. The list of these genes illuminates distinct physiological roles of this organ. A number of genes are related to extracellular matrix (ECM), including KH.C1.1211 (laminin), KH.C5.562 (dystroglycan), KH.C8.476 (lysyl oxidase like 3), KH.C9.371 (discoidin domain receptor tyrosine kinase 2), KH.C13.31 (secreted protein acidic and cysteine rich), KH.L87.44 (versican), KH.L139.15 (hemicentin-1), KH.C13.129 (sushi, von Willebrand factor type A, EGF and pentraxin domain containing 1), KH.C1.707 (mucin 13), and KH.C1.1291 (matrix metallopeptidase 16). Another conspicuous feature is the expression of prospective neural support and axon guidance molecules, such as KH.C12.72 (netrin 1), KH.C5.524 (netrin 4), KH.C5.490 (netrin receptor unc-5), KH.L18.70 (neuregulin 2), KH.C11.251 (nerve growth factor receptor), and KH.S749.1 (glial fibrillary acidic protein, GFAP). Transmembrane receptors and adhesion molecules are also noticeable, including KH.C11.90/KH.C8.337 (somatostatin receptors), KH.C2.341 (G-protein-coupled receptor 107), KH.C14.202 (insulin receptor related receptor), KH.C9.149 (prostaglandin I2 receptor), KH.C9.260 (frizzled class receptor 5/8), KH.C14.562 (adhesion GPCR D1), KH.C14.88 (FAT atypical cadherin 1), and KH.C12.14 (cadherin 18). The marker genes for the neural gland also include genes for solute carrier family (SLC) transporters (KH.C7.199, KH.L4.19, KH.S215.18), calponin (KH.C4.559), and proteins involved in cell growth and differentiation, such as KH.C2.605 (hedgehog interacting protein), KH.C2.957 (msh homeobox), KH.C4.236 (Elf3/Ehf transcription factor), KH.C7.22 (ectodysplasin A), and KH.L112.16 (Jun/AP-1 transcription factor subunit). The coordinated enrichment of these molecules suggests distinct cell–cell interaction and extracellular environment properties within the neural gland, consistent with its proposed secretory and interface-related roles.

### 5. Identification of new marker genes for the cerebral ganglion

We next focused on the cerebral ganglion. The cerebral ganglion is divided into the cortex and medulla ^17,18^. The super-resolution gene expression maps revealed that spatial expression patterns could be broadly classified into four major groups: i) uniformly expressed in the cortex (cortex), ii) predominantly expressed in the central cortex region, particularly in the ventral region facing the neural gland (central), iii) predominantly expressed in anterior and posterior parts of the cortex (anteroposterior), and iv) predominantly expressed in the inner medulla region (medulla). For the cortex (**Fig. 3D; Supplementary Table S2**), we identified marker genes including KH.C7.92 (NMDA receptor), KH.C12.16 (carboxypeptidase E), KH.L128.2 (proprotein convertase 2), KH.S761.6 (glutamate decarboxylase), KH.C6.11 (aminopeptidase O), KH.S618.6 (galanin-like neuropeptide precursor), KH.C8.904 (Dickkopf 3) and KH.C1.215 (acid sensing ion channel subunit 1/2/3). Many of these genes are associated with the biogenesis of gamma-aminobutyric acid (GABA) and neuropeptides, and synaptic function and proton-gated ion channel activity, suggesting active neuronal signaling and peptide processing activities in this region. For the central cortex domain (**Fig. 3E; Supplementary Table S3**), we identified marker genes including KH.C1.421 (4-aminobutyrate aminotransferase, ABAT), KH.C1.498 (choline O-acetyltransferase), KH.C12.218 (acetylcholine esterase), KH.C9.885 (vasopressin receptor), KH.C1.1067 (neuronal calcium sensor 1), KH.C2.101 (synaptotagmin), KH.C5.389 (tachykinin precursor), KH.L22.28 (type B GABA receptor), KH.C1.467 (μ-opioid/somatostatin/angiotensin receptor-like), KH.S1051.1/KH.S1104.4 (gonadotropin-releasing hormone, GnRH, precursor) and KH.C9.462 (voltage-gated Na^+^ channel). These genes suggest specialized neurotransmitter response and synaptic vesicle release properties in this subregion. For the anteroposterior distal regions (**Fig. 3F; Supplementary Table S4)**, we identified marker genes including KH.C2.491 (golgin A4), KH.C4.437 (Ca^2+^/calmodulin-dependent protein kinase IV), KH.C6.106 (phosphodiesterase 4D interacting protein), KH.C3.474 (MAM domain containing glycosylphosphatidylinositol anchor 2; MDGA2), and KH.C7.399 (calcitonin). For the medulla (**Fig. 3G; Supplementary Table S5**), we identified marker genes including KH.L55.2 and KH.C10.335 (calmodulin family members), KH.C9.150 (doublecortin-like and CAM kinase-like 2), KH.C8.892 (α-tubulin), KH.L116.85 (β-tubulin), and KH.C1.342 (voltage-gated Ca^2+^ channel). These genes are related to calcium signaling and cytoskeletal structure, consistent with structural core and *regulatory function* in the cerebral ganglion.

## DISCUSSION

In this study, we present a spatially resolved transcriptomic atlas of the adult *Ciona* neural complex. Clustering-based analysis defined the major tissue domains of the adult neural complex, providing a comprehensive transcriptional framework for understanding its organization and surrounding structures. In addition to the cerebral ganglion and neural gland, the atlas also resolves adjacent tissues such as the ciliated funnel, the neural gland duct, and the dorsal strand, offering a spatial molecular resource that may facilitate future studies of these structures.

To further refine spatial resolution in adult tissues, we generated super-resolution gene expression maps by integrating histological image features with spatial transcriptomic data. This approach provides a scalable molecular reference that reduces reliance on gene-by-gene anatomical screening. Based on the super-resolution gene expression maps, we identified spatially restricted marker genes for both the neural gland and the cerebral ganglion. The neural gland displays a distinct molecular signature consistent with specialized structural and secretory functions, while the cerebral ganglion exhibits clear internal zonation, indicating functional compartmentalization within this compact brain. These findings suggest that the spatial organization of neuronal and neuroendocrine programs is already established in a simple chordate nervous system. Previous studies have identified conserved molecular signatures shared between *Ciona* and vertebrates in particular cell types, including photoreceptor-related cells ^19^ and hypothalamic neuroendocrine lineages ^2^. The atlas presented here extends these efforts by providing a comprehensive spatial framework for comparing conserved molecular signatures across species, thereby enabling systematic investigation of how regional specialization of the vertebrate brain emerged during evolution.

Special attention has been paid to the neural gland due to its distinct anatomical features and possible homology to vertebrate organs. The proposed homologous organs in vertebrates include adenophypophysis ^20,21^ and choroid plexus ^16^. Various functions have been proposed for the neural gland, including an endocrine organ ^21–23^, an excretory organ ^24^, an exocrine organ ^16^, a lymph producing organ ^22^, an osomoregulatory organ ^16,25^, and an organ that mediates regeneration of the cerebral ganglion ^26,27^. Despite these interests and studies, the physiological and developmental roles of this organ remain elusive. Accordingly, only a few genes have been identified as specifically expressed in the neural gland ^8,16^.

In the present study, we identified 85 neural gland-specific genes. These genes support some of the previously proposed functions and homology of the neural gland. Expression of various SLC transporters with calponin suggests its role in regulating chemical (ions, amino acids, and peptides) and physical (pressure or flow of fluid) environment around the neural complex, or acting as a selective barrier, similar to a blood-brain barrier of vertebrates. A osmoregulatory role is also suggested by the expression of various GPCRs for the putative osmoregulatory neuropeptides; some of these receptors are also expressed in the vertebrate choroid plexus ^16^. Deyts et al. (2006) proposed that the neural gland is an osmoregulatory organ and might be homologous to the choroid plexus of vertebrates. Furthermore, expression of GFAP and neural guidance cues (netrins, netrin receptors, and neuregulin 2) supports a proposed role of the neural gland in development and regeneration of the cerebral ganglion. Our results are compatible with this idea and further extend its neuroprotective and supportive roles, similar to the blood-brain barrier of vertebrates. The richness of ECM proteins and putative protective and instructive roles for the brain are also reminiscent of the meninges of vertebrates, especially the pia mater, where the choroid plexus derive from. Thus, the neural gland may represent a primitive form of the meninges-choroid plexus system in vertebrates.

Previously, the cortex and medulla are recognized as distinct regions in the cerebral ganglion ^17,18^. Our super-resolved gene maps identified four groups of gene expression patterns in the cerebral ganglion: i) uniformly expressed in the cortex, ii) predominantly expressed in the central cortex region, particularly in the cortex region facing the neural gland, iii) predominantly expressed in the anterior and posterior parts of the cortex, and iv) predominantly expressed in the medulla. Pan-cortex and central cortex genes include genes for the biogenesis of neurotransmitters and neuropeptides, their receptors, and various proteins involved in synaptic function, suggesting that the cortex contains a number of neurons and serves as the primary site of neuronal activity. On the contrary, the anteroposterior distal regions of the cortex show a distinct molecular signature in which proteins associated with neuronal and synaptic functions are less prominent, suggesting a distinct molecular function of these regions. In the medulla, genes for proteins involved in intracellular Ca^2+^ signaling are conspicuously expressed together with microtubule components. Expression of these genes suggests the medulla serves as the structural core of the cerebral ganglion and plays a regulatory role with active Ca^2+^ signaling, which is reminiscent of astrocytes in the vertebrate brain. In summary, the present analysis revealed previously unrecognized molecular heterogeneity in the cerebral ganglion, especially within the cortex.

Several limitations should be acknowledged. First, the current spatial maps are derived from spot-based measurements, and integration with single-cell RNA sequencing data from the adult brain will be necessary to further refine cell-type–specific expression patterns. Such integration would enable more precise delineation of cellular heterogeneity within each anatomical domain ^28^. Second, we did not perform extensive regulatory network analysis. In *Ciona*, widespread trans-splicing complicates transcript-level assignment and regulatory inference ^29^, posing challenges for conventional gene regulatory network reconstruction. Future studies combining improved transcript annotation, single-cell data, and perturbation experiments will be essential for dissecting regulatory mechanisms underlying the spatial programs described here.

## DATA AVAILABILITY

FASTQ data from this study can be downloaded from GEO database (GSE327796). Processed data can be downloaded from figshare (https://doi.org/10.6084/m9.figshare.31915545). Codes generated for this project are available on GitHub (https://github.com/xzengComBio/Ciona_ST).

## AUTHOR CONTRIBUTIONS

Conceptualization, X.Z., F.G., T.G.K., and K.N.; Visium Data Acquisition, A.M., N.O., K.M., T.G.K., and Y.S.; Data Curation, X.Z. and T.G.K.; Formal Analysis, X.Z.; Investigation, X.Z., F.G., A.M., and T.G.K.; Writing – Original Draft, X.Z., F.G., and T.G.K.; Writing – Review & Editing, X.Z., F.G., T.G.K., and K.N.; Funding Acquisition, T.G.K. Y.S., and K.N.; Resources, K.M., Y.S., T.G.K., and K.N.; Supervision, T.G.K. and K.N.

## ACKNOWLEDGEMENTS

We thank the National BioResource Project of MEXT and all members of the Maizuru Fisheries Research Station and Yutaka Satou Lab of Kyoto University and the Misaki Marine Biological Station and Manabu Yoshida Lab of the University of Tokyo for providing us with C. intestinalis type A adults. We also thank the Graduate School of Maritime Sciences, Kobe University, for generously allowing us to collect C. intestinalis type A on the campus. Computational resources were provided by the supercomputer system SHIROKANE at the Human Genome Center, Institute of Medical Science, the University of Tokyo.

## MATERIALS AND METHODS

### Biological materials

Mature adults of *Ciona intestinalis* type A were provided by the Maizuru Fisheries Research Station of Kyoto University and by the Misaki Marine Biological Station of the University of Tokyo through the National Bio Resource Project (NBRP) of the Ministry of Education, Culture, Sports, Science and Technology of Japan (MEXT). These were maintained in indoor tanks of artificial seawater (ASW) (Marine Art BR; Tomita Pharmaceutical, Tokushima, Japan) at 17-20°C.

### Tissue preparation and VISIUM sequencing library construction

*Ciona* whole brains were surgically dissected from mature adult individuals, embedded in Tissue-Tek OCT compound (Sakura Finetek, Tokyo, Japan), and sectioned at 10-µm thickness using a cryostat (HM525, Microm). Tissue fixation and staining were as described in the Methanol Fixation, H&E Staining & Imaging for Visium Spatial Protocols (10X Genomics; #CG000160 Rev C). Tissue permeabilization optimization was performed using Visium Spatial Tissue Optimization Slide & Reagent Kit (10□×□Genomics, #1000193) according to the Visium Spatial Tissue Optimization User Guide Rev B (10□×□Genomics, #CG000238). The optimal tissue permeabilization time for *Ciona* brains was determined as 12 min. The optimized condition was then used for Visium Spatial Gene Expression library preparation. The four capture areas on the Visium Gene Expression Slide (#2,000,233) contained serial sections from an embedded brain block, including three brains from different individuals. When embedded, the brain tissues were aligned in the same direction so that each capture area contained three sagittal brain sections. Each capture area of the Visium Gene Expression Slide contained approximately 5000 barcoded spots of 55 μm in diameter (100-μm center-to-center spacing between spots). The sequencing library was prepared using Visium Spatial Gene Expression Slide & Reagent Kit (10□×□Genomics, #1000184) according to the Visium Spatial Gene Expression User Guide Rev C (10X Genomics, #CG000239) and sequenced on a NovaSeq 6000 system (Illumina). Sequencing was performed with the recommended protocol (read 1: 28 cycles; i7 index read: 10 cycles; i5 index read: 10 cycles; and read 2: 90 cycles), yielding between 391 million and 440 million sequenced reads.

### Mapping

Sequencing output was processed through the Space Ranger 1.3.1 count function using default parameters. Reads were quantified using the KH2013 reference genome provided by the Ghost Database ^30^. The raw count matrices were subsequently imported into R and subjected to analysis using the Seurat 4 package ^15^. The data were log normalized and scaled by the SCTransform function ^31^, followed by batch effect correction using canonical correlation analysis. The corrected data were scaled for the principal component analysis (PCA). The corrected data was used for clustering based on a shared nearest-neighbor modularity optimization-based clustering algorithm with default parameters and resolutions at 0.2. Non-linear dimensionality reduction was conducted using UMAP.

We performed Gene Ontology enrichment analysis for cluster-specific marker genes identified from spatial transcriptomic data using a custom *Ciona* GO annotation derived from the Aniseed GAF file ^32^. Enrichment of Biological Process terms was tested with clusterProfiler ^33^, and the top-enriched GO terms for each cluster were visualized using dot plots.

### Generation of super-resolution gene maps

We used xfuse^14^ to generate high resolution gene expression map for marker genes. First, the coordinates on each brain section were extracted from the original “tissue_positions_list.csv” file. Then, the transcriptomics and image information for each brain section were extracted using xfuse convert Visium function with a scale value setting to 0.2. Finally, the separated 12 transcriptomics and image section were train by the xfuse model using the xfuse run function via Nvidia V100 GPU. The convergence of the model was achieved following a computational duration of approximately 7 hours.

