## supplementary figures for "Spatial transcriptomic landscape of the *Ciona* adult brain: functional zonalisation and cellular composition in a sessile chordate brain and a novel insight into neural gland function"

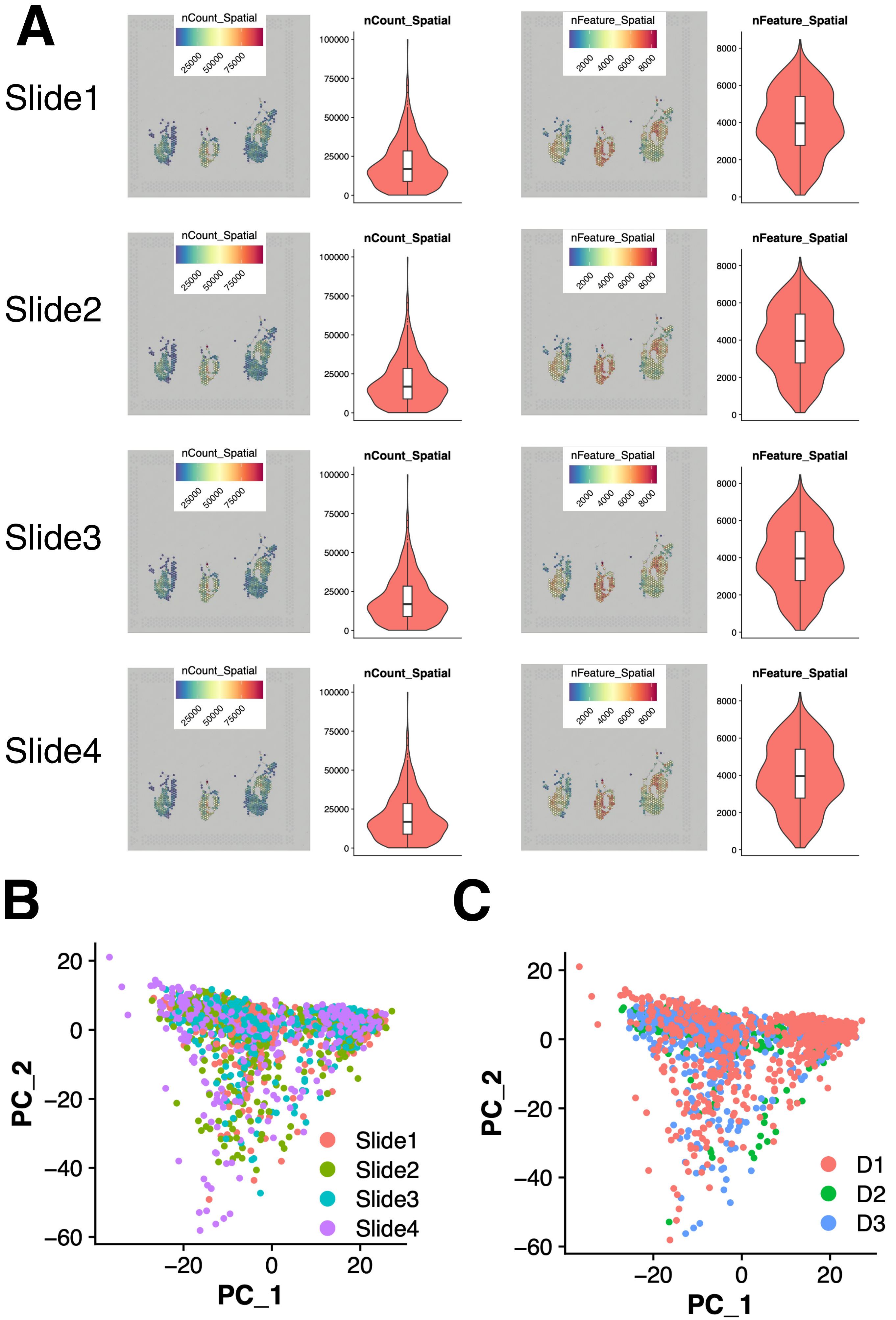
A
Slide1
Slide2
Slide3
Slide4
nCount_Spatial
25000
50000
75000
nFeature_Spatial
2000
4000
6000
8000
100000
0
B
C
PC_1
PC_2
Slide1
Slide2
Slide3
Slide4
D1
D2
D3

**Supplementary Figure 1. Quality control and sample integration of Visium spatial transcriptomic datasets.**

**A** Quality control metrics for four Visium slides. For each slide (Slide1–Slide4), spatial distributions of total UMI counts per spot (nCount_Spatial) and detected gene numbers per spot (nFeature_Spatial) are shown. Corresponding violin plots summarize the distribution of these metrics across all spots under tissue for each slide. **B-C** Principal component analysis (PCA) of all retained spots colored by slide identity (**B**) and biological donor (D1–D3) (**C**) after SCTransform normalization and CCA-based integration.

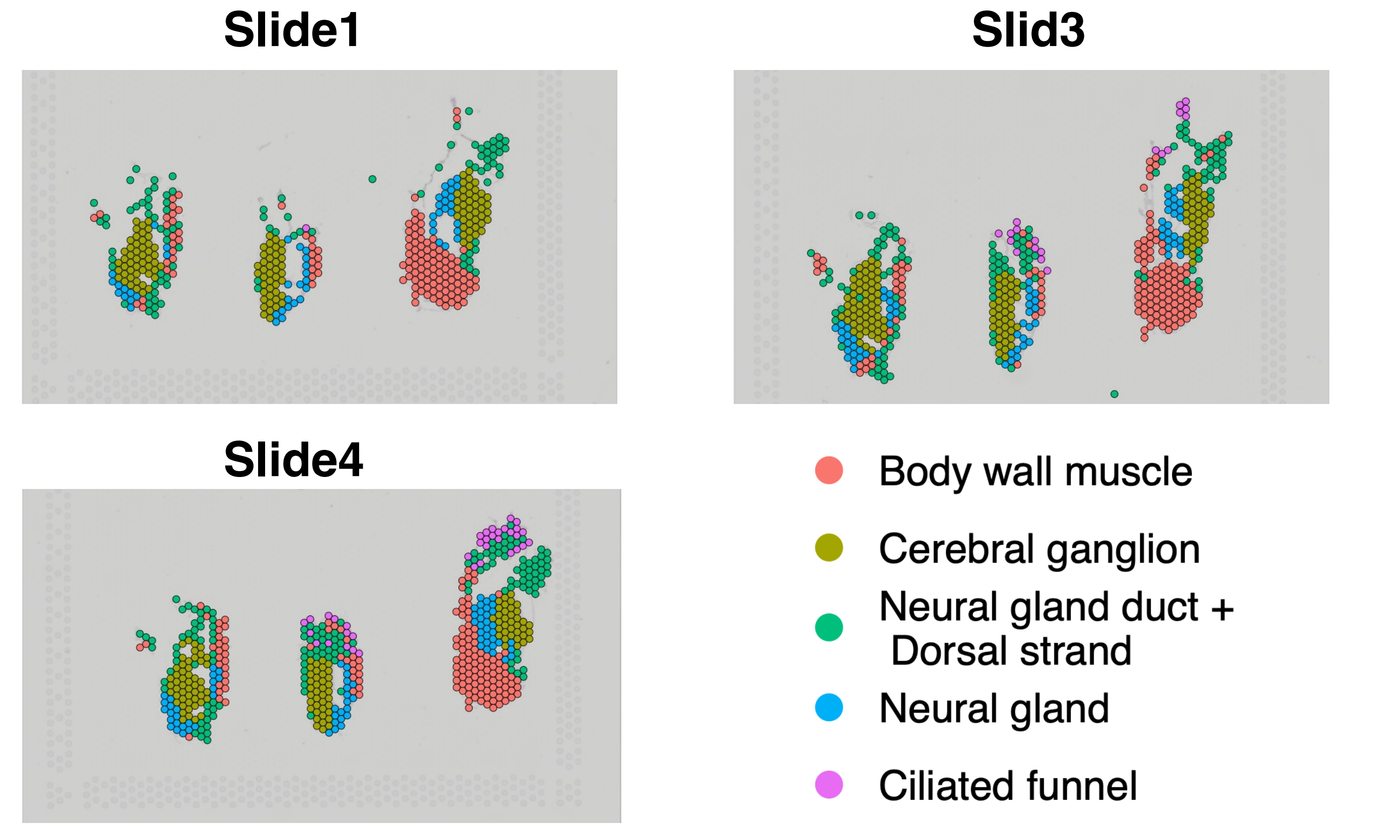


**Supplementary Figure 2. Reproducibility of spatial clustering and functional annotation of identified tissue domains.**
Spatial distribution of clusters across representative Visium slides (Slide1, Slide3, and Slide4).

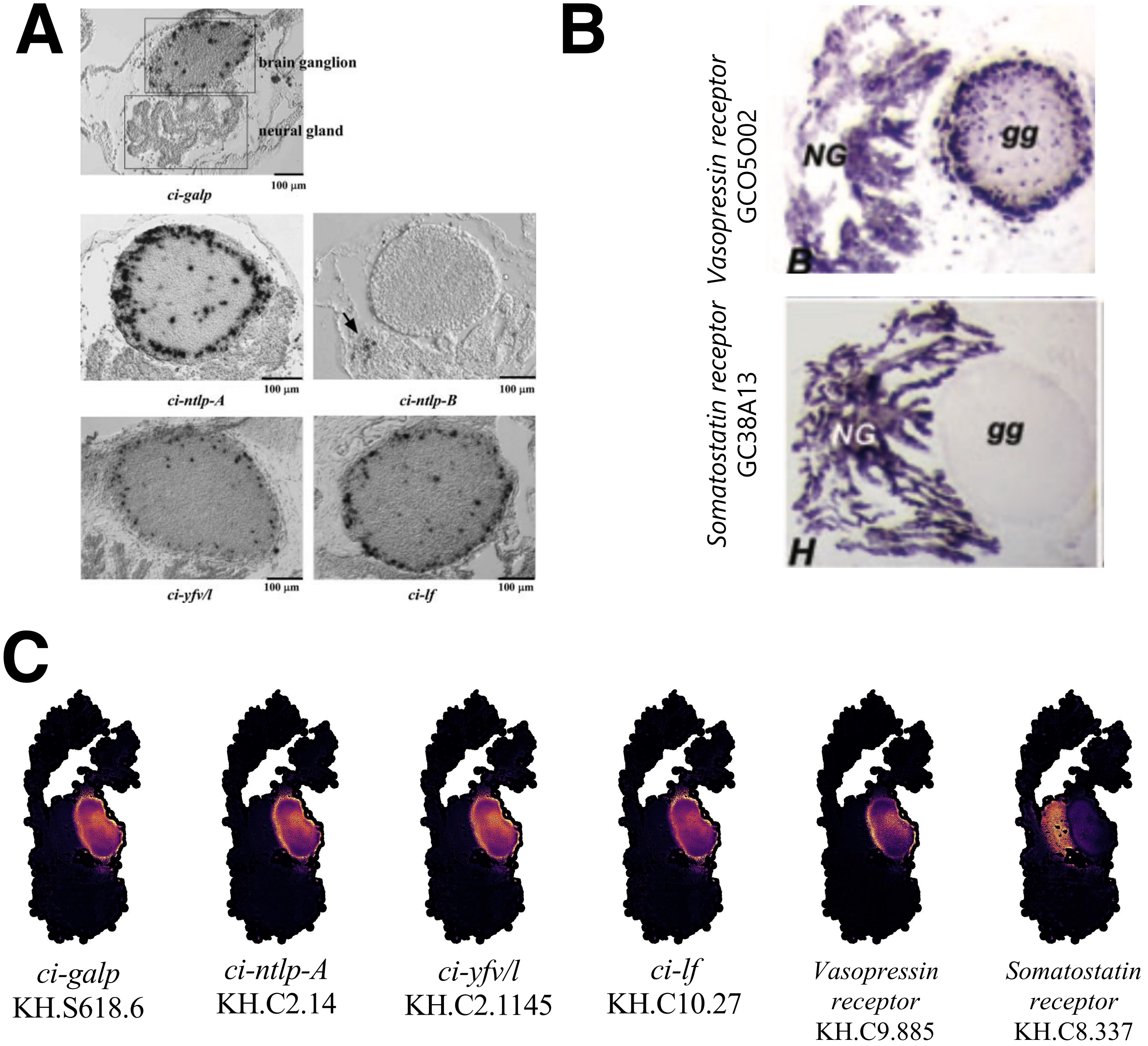


**Supplementary Figure 3. Independent validation of spatial expression patterns by in situ hybridization.**

**A** Representative in situ hybridization images showing expression patterns of *ci-galp*, *ci-ntlp-A*, *ci-ntlp-B*, *ci-vfyl/l*, and *ci-lf* in transverse sections of the adult neural complex (from Kwada et al. 2011). **B** Representative in situ hybridization images showing expression patterns of *vasopressin receptor* and *somatostatin receptor* (from Deyts et al. 2006) **C** Corresponding spatial transcriptomic expression maps for the same genes generated from Visium data (super-resolution reconstruction).
